# Evolutionary genomics of two co-occurring congeneric fore reef coral species on Guam (Mariana Islands)

**DOI:** 10.1101/2024.10.15.618583

**Authors:** Héctor Torrado, Dareon Rios, Karim Primov, David R Burdick, Bastian Bentlage, Sarah Lemer, David Combosch

## Abstract

Population structure provides essential information for developing meaningful conservation plans. This is especially important in remote places, such as oceanic islands, where limited population sizes and genetic isolation can make populations more susceptible and self-dependent.

In this study, we assess and compare the relatedness, population genetics and molecular ecology of two sympatric *Acropora* species, *A. surculosa sensu* Randall & Myers (1983) and *A.* cf. *verweyi* Veron & Wallace, 1984 around Guam, using genome-wide sequence data (ddRAD). We further contrast our findings with the results of a recent study on back reef *A.* cf. *pulchra* (Brook, 1891) to assess the impact of habitat, colony morphology and phylogenetic relatedness on these basic population genetic characteristics and generate testable hypotheses for future studies.

Both target species were found to have small effective population sizes, low levels of genetic diversity, and minimal population structure around Guam. Nonetheless, *A.* cf. *verweyi* had significantly higher levels of genetic diversity, some population structure as well as more clones, close relatives and putative loci under selection. Comparisons with *A.* cf. *pulchra* indicate a potentially significant impact by habitat on population structure and genetic diversity while colony morphology seems to significantly impact clonality.

This study revealed significant differences in the basic population genetic makeup of two sympatric *Acropora* species on Guam. Our results suggest that colony morphology and habitat/ecology may have a significant impact on the population genetic make-up in reef corals, which could offer valuable insights for future management decisions in the absence of genetic data.

## Introduction

Studying and maintaining population connectivity is critical for coral reef conservation since it significantly enhances chances for faster reef recovery (Almany et al. 2009; Cros et al. 2017; Jones et al. 2009). Among reef-building corals, species from the *Acropora* genus hold significant importance, with distributions in coral reefs around the world, from the Red Sea, to the Indo-Pacific Ocean and the Caribbean. It represents the most diverse and abundant coral taxon, with over 100 described species (Wallace, 1999), which is currently being revised and will likely increase significantly in the near future (e.g., Bridge et al. 2023; Cowman et al. 2020). *Acropora* corals occur across a wide variety of habitats (intertidal, lagoons, reef slopes etc.) and with a diverse range of morphologies (e.g. arborescent, hispidose, corymbose, table) (Wallace 1999). Their rapid growth rates contribute significantly to reef accretion, which allow them to rapidly colonize new habitats or recover areas following mortality events (e.g. Tomascik et al. 1996). The complex three-dimensional structures created by *Acropora* corals provide refuge for a myriad of marine organisms, making them crucial for habitat preservation and reef ecology (Knowlton et al. 2010). However, a substantial proportion of *Acropora* species are highly susceptible to bleaching caused by rising sea surface temperatures, with over 70% listed as near threatened or threatened by the International Union for Conservation of Nature Red List (Carpenter et al. 2008).

In this study, we focus on two distinct *Acropora* species, *A.* cf. *verweyi* Veron & Wallace (1984) and *A. surculosa sensu* Randall & Myers (1983). With the *Acropora verweyi* Veron & Wallace, 1984 type locality in the Cocos-Keeling Islands in the Eastern Indian Ocean, it is likely that the species referred to in the Mariana Islands as *A. verweyi* represents a genetically distinct, but morphological similar, taxon. As such, we apply the Open Nomenclature qualifier “cf.” when referring to this taxon. *Acropora surculosa* was first described by Dana (1846) from Fiji and first recorded as such in Guam and the Marianas Islands by the late Richard Randall in Marine Laboratory (1981) and pictured in the Randall & Myers (1983) field guide. However, Veron & Wallace (1984) later synonymized Dana’s *A. surculosa* with *A. hyacinthus*, it was retained as a junior synonym of *A. hyacinthus* in the seminal treatise of the genus by Wallace (1999) and is currently recognized as *A. hyacinthus* (Hoeksema & Cairns 2024). Subsequent molecular studies of *A. hyacinthus* revealed a species complex with multiple cryptic species, including two cryptic species in Pohnpei and three in Palau (Ladner & Palumbi 2012). Some of these cryptic species were later identified in other locations, including American Samoa (Rose et al. 2021), Japan and Taiwan (Suzuki et al. 2016). A recent taxonomic revision of the *Acropora hyacinthus* complex (Fitzgerald et al., in prep) found that *A. hyacinthus* (Dana, 1846) was a distinct lineage from *A. surculosa sensu* Randall & Myers (1983), and are in the process of describing the latter as a new species, with Guam as a the type locality (Bridge, pers. comm.). Since these results have yet to be published, our use of the name *Acropora surculosa* henceforth is specifically in reference to *A. surculosa sensu* Randall & Myers (1983) to avoid confusion with other lineages in the *A. hyacinthus* species complex and facilitate the correspondence of our data to the putatively new species.

The two species differ in several fundamental morphological and life history traits. For example, *Acropora surculosa* forms corymbose colonies with robust, tapering branches while *A.* cf. *verweyi* is digitate or cushion-shaped with more delicate, thinner, terete branches. Additionally, *Acropora* cf. *verweyi* is more bleaching susceptible than *A. surculosa sensu* Randall & Myers (1983) (Maynard et al. 2018), although both species occur in similar habitats on upper reef slopes (Veron, 2000). They also have a potentially similar Indo-Pacific-wide distribution if *A. surculosa* is considered synonymous to *A. hyacinthus* (Veron, 2000) - otherwise, the distribution range of the Marianas *A. surculosa sensu* Randall & Myers (1983) is unknown, but likely much smaller.

While some *Acropora* species have received considerable attention, the majority of population genomics studies have been limited to a handful of species, mostly in the Caribbean (e.g. Drury et al., 2016), the Ryukyu Islands (e.g. Selmoni et al., 2020; Shinzato et al., 2015), and the Great Barrier Reef (e.g. Fuller et al., 2020). Beyond these focus areas, there is a general scarcity of studies, even though remote oceanic islands are an important habitat for these corals (but see Cros et al. 2017; Davies et al. 2015). This knowledge gap poses challenges for planning and implementing effective conservation strategies in regions primarily composed of such habitats, such as Guam in the Mariana Islands and the broader Micronesian region.

Guam is the largest and southernmost island in the Marianas archipelago in Micronesia. Reefs in this island experienced substantial coral losses over a five-year period (2013–2017) due to consecutive bleaching events and extreme low tides (Raymundo et al. 2017, 2019; Reynolds et al. 2014). Consequently, high mortality rates may have impacted the population structure and diversity, potentially generating bottlenecks in the coral population on Guam. This situation makes the study of Guam particularly intriguing, especially considering the ongoing development of diverse conservation and restoration plans on the island (e.g. Raymundo et al. 2022).

In this study, we assessed the genetic connectivity, population structure and genetic diversity of two congeneric and sympatric coral species on Guam: *A. surculosa* and *A.* cf. *verweyi*. One goal was to assess if there are fundamental differences in the population genetic setup between a more bleaching susceptible and a more resilient species, especially in terms of clonality and local adaptations. We employed a ddRAD-Seq approach to generate genome-wide sequencing data for 2 species and 3 Guam populations each. By comparing the results for these two phylogenetically distinct but ecologically similar species with recent results for another *Acropora* species, *A.* cf. *pulchra* (Rios et al. 2024), we shed light on the patterns, underlying causes, and ecological consequences of intraspecific and interspecific population genomic variation among these coral populations, inhabiting small oceanic islands.

## Results

On average, 535,799 reads were obtained per *Acropora* cf. *verweyi* sample, which resulted in 213,507 STACKS loci. For *A. surculosa sensu* Randall & Myers (1983), on average 214,507 reads and 46,494 STACKS loci were obtained per sample. After filtering and data curation, we kept 1,642 loci and 51 *A.* cf. *verweyi* samples as well as 1,168 loci and 38 *A. surculosa* samples. These loci were genotyped in at least 50% of samples per species, with a mean depth per locus of 33.0 ± 8.8 and 31.8 ± 6.3 reads, respectively. Sample and loci filters significantly reduced the *A. surculosa* dataset and, for example, excluded all samples from the Ritidian population, while only one *A.* cf. *verweyi* sample was removed (Figure 1, Table 1).

**Fig. 1.**
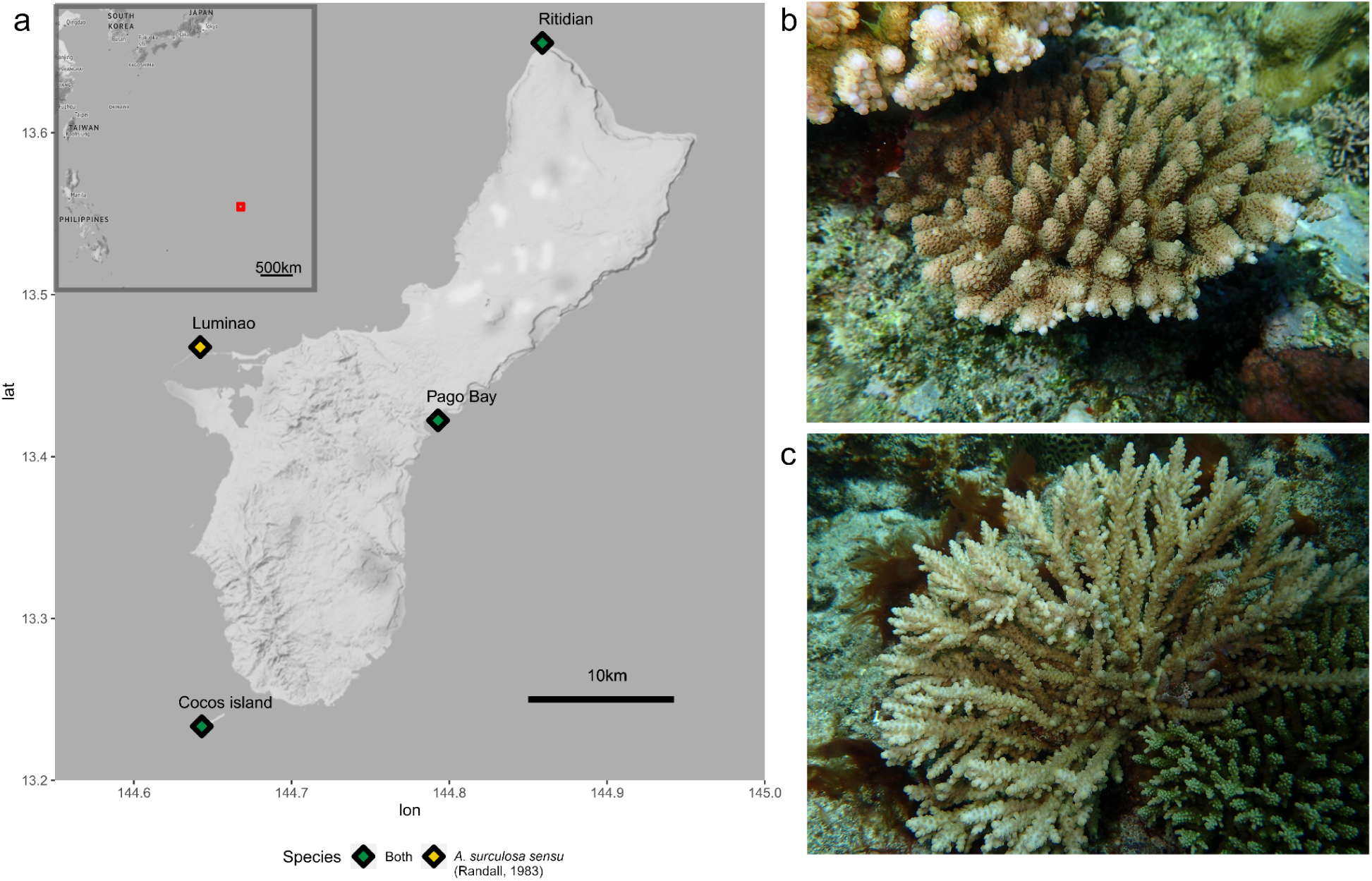
Map of the sampling locations (diamonds) [a] and species pictures [b, c]. a) Map. The red square on the big scale map corresponds to the enlarged map of the analyzed area. Sampling map was created using ggmap (Kahle & Wickham 2013) and ggplot2 (Wickham, 2009). b) *Acropora surculosa sensu* Randall & Myers (1983) was sampled from all four sites c) *A.* cf. *verweyi* was sampled in 3 of the 4 sites but was temporarily extirpated and thus not sampled at Luminao.

**Table 1.** Overall population genetic stats for both species.

| Species | N | N<sub>g</sub> | N<sub>g</sub>/N | SNPs | H<sub>O</sub>- SNP<br>(95% CI) | H<sub>E</sub>- SNP<br>(95% CI) | G<sub>IS</sub>- SNP<br>(95% CI) | N<sub>e</sub><br>(95% CI) |
| --- | --- | --- | --- | --- | --- | --- | --- | --- |
| *A. verweyi* | 51 | 41 | 0.8 | 1,642 | 0.239<br>(0.230-0.249) | 0.250<br>(0.242-0.258) | 0.043<br>(0.014-0.053) | 670<br>(548-1110) |
| *A. surculosa* | 36 | 36 | 1.0 | 1,168 | 0.127<br>(0.119-0.135) | 0.154<br>(0.146-0.163) | 0.178<br>(0.143-0.191) | 247<br>(191-337) |

Phylogenetic analyses revealed that our two target species are very distantly related to each other (Figure 2): *Acropora* cf. *verweyi* is most closely related to a clade composed of *A. tenuis* and *A. yongei*, which is sister to all other *Acropora* species included in this analysis. In contrast, *A surculosa* formed a clade with *A. hyacinthus* from Okinawa, which corresponds to its currently valid species name (*sensu* Veron & Wallace (1984), see introduction and discussion for further information about this topic). This clade was most closely related to *A. cytherea* (Dana, 1846) from Okinawa. The third species from Guam, *A.* cf. *pulchra*, formed a clade with *A. millepora* from the GBR (Fuller et al 2020), which was closely related to *A. selago* (Studer 1879). *A.* cf. *pulchra* appears to be more closely related to *A. surculosa* than to *A.* cf. *verweyi* (Figure 2).

**Fig. 2.**
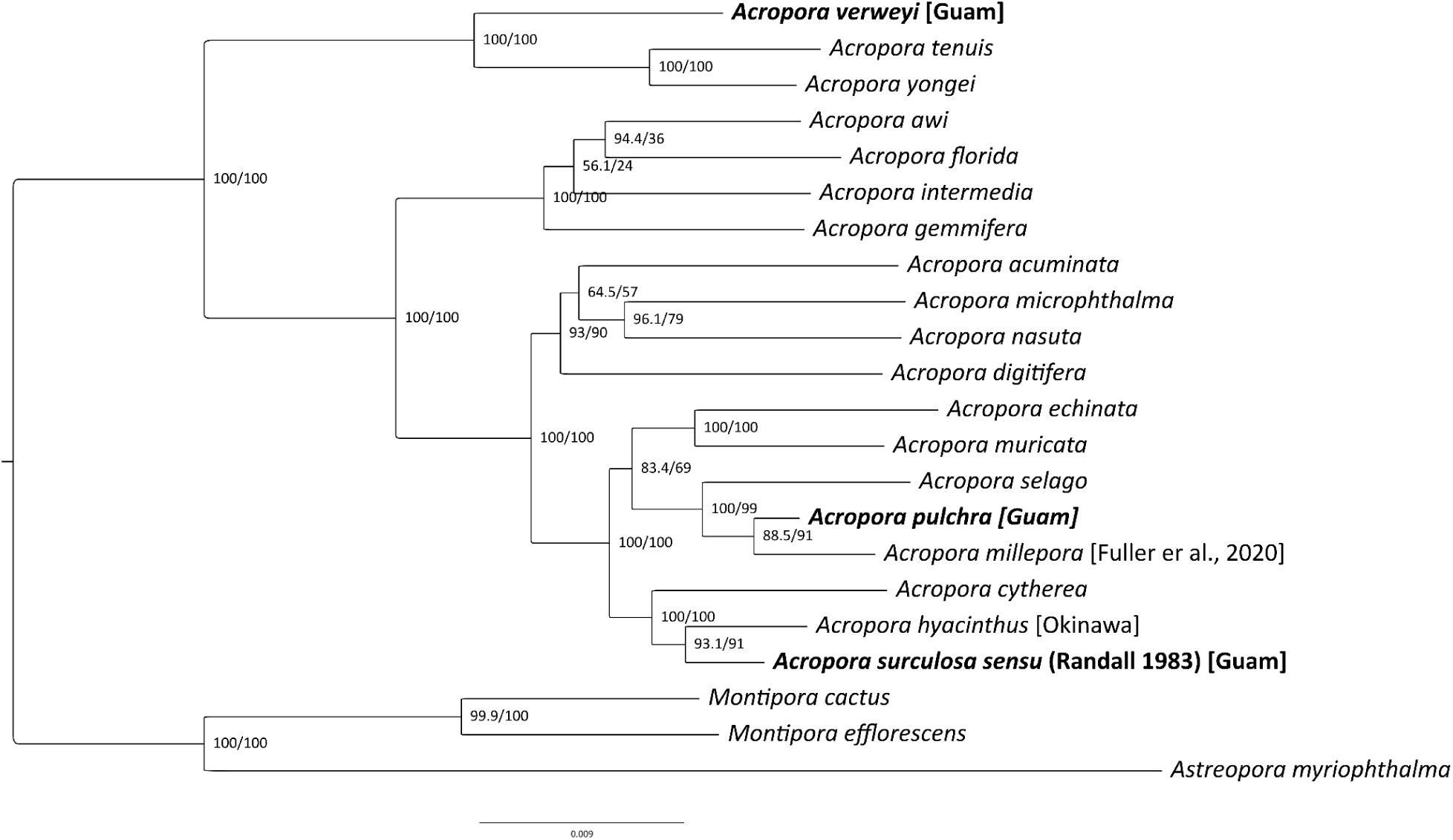
Maximum likelihood phylogenetic tree for *Acropora* species based on 50,583 bp over 52 loci. Numbers in nodes represent SH-aLRT (%) and ultrafast bootstrap support (%). Samples from Guam are in bold.

IBS-based cladograms did not reveal any outliers, which confirms that all samples belong to just two species (Figure S1). Cutoff values to distinguish sexually from asexually generated genotypes were determined by the first large gap among IBS-based distances, which was determined to be 0.08 for *A.* cf. *verweyi* and 0.04 for *A. surculosa*. For *A.* cf. *verweyi,* 8 pairs and 1 triplet of putative clones were identified. In contrast, no clones were identified for *A. surculosa*. After removing technical replicates and leaving only one ramet per genet, 41 individuals of *A.* cf. *verweyi* and 36 individuals of *A. surculosa* were retained (Table S1). The number of clones was significantly higher in *A.* cf. *verweyi* (Fisher’s exact test, p-value = 0.0045).

Both species show no significant small-scale spatial genetic structure between 0 and 200m in the genet datasets, i.e. there is no clear pattern of elevated relatedness over small spatial scales among sexually derived genotypes (Figure S2). In contrast, in the ramet dataset, there is a significant spatial genetic structure over the first 20 m for *A.* cf. *verweyi,* i.e. the observed average relatedness for individuals 20 m apart is beyond the 95% confidence intervals. In other words, clones significantly elevate the average relatedness among nearby but not distant samples. This difference is clearly indicated by the Sp statistic as well, which is −0.012 for *A. surculosa sensu* Randall & Myers (1983), −0.001 for *A.* cf. *verweyi* and 0.018 for *A.* cf. *verweyi* including clones.

Genetic diversity was low in both species, especially in *A. surculosa* (Table 1). For both species, the observed heterozygosity is slightly smaller than the expected heterozygosity. The difference is more pronounced in *A. surculosa*, leading to a higher heterozygote deficit (Table 1, Table S2), with non-overlapping 95% confidence intervals (CI) for the two species, which we interpret as significantly different. In line with these results, the effective population size (based on linkage disequilibrium) is small for both species, but almost three times bigger for *A.* cf. *verweyi* than for *A. surculosa* (670 vs 247; Table 1) - also with non-overlapping CI.

The site frequency spectrum (SFS) had a higher abundance of alleles of low frequency class in *A. surculosa* while *A.* cf. *verweyi* had a higher abundance of high frequency class sites in both the genome-wide and SNP-based datasets (Figure S4). Genome-wide Tajima’s D values were negative for both species in the genome-wide dataset, with more negative values in *A. surculosa*, with an average of −0.512 (± 0.819), compared to −0.103 (± 1.106) for *A.* cf. *verweyi* (Figure S5). An excess of rare, low-frequency alleles and the more negative Tajima D values in *A. surculosa* both suggest a recent population expansion, compared to *A.* cf. *verweyi*.

Looking at population structure, AMOVA results for both species show that the vast majority of genetic variation was partitioned within rather than between locations in both species (Table 2a). The proportion of genetic variation partitioned by populations was insignificant (p = 0.55) in *A. surculosa*, which aligns with the absence of any obvious population structure in the DAPC (Figure 3a) and ADMIXTURE graphs (Figure S3b). In contrast, in *A.* cf. *verweyi*, a small but significant proportion of the total genetic variation was partitioned between populations (1.5%; p = 0.01). This difference is not very obvious in the DAPC but in comparison with *A. surculosa*, *A.* cf. *verweyi* populations are more distinct and more clearly and cleanly separated (Figure 3). The ADMIXTURE results do not show any obvious population structure for *A.* cf. *verweyi* (Figure S3).

**Fig. 3.**
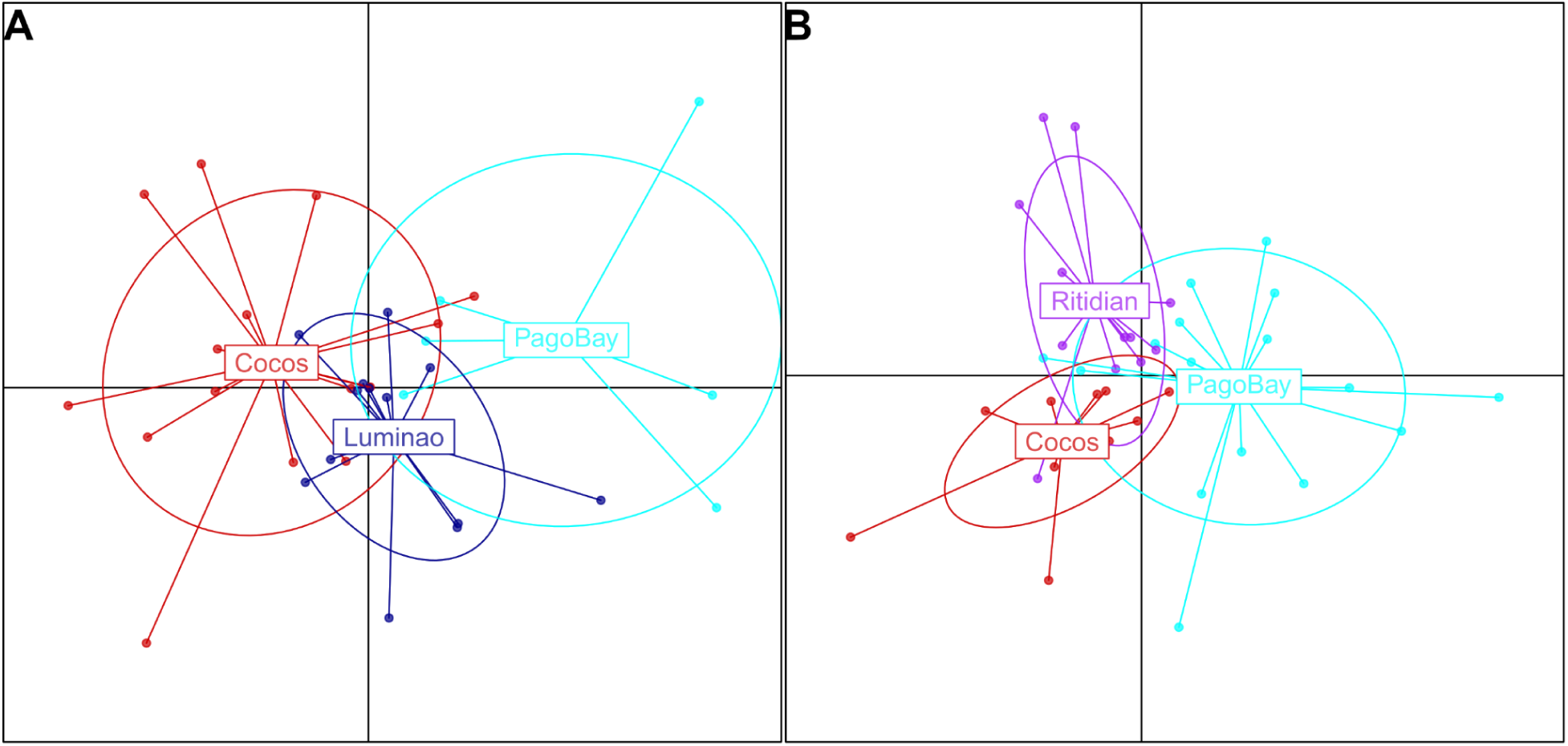
Discriminant Analysis of Principal Components (DAPC) results representation for A) *A. surculosa sensu* Randall & Myers (1983) and B) *A.* cf. *verweyi*.

**Table 2:**
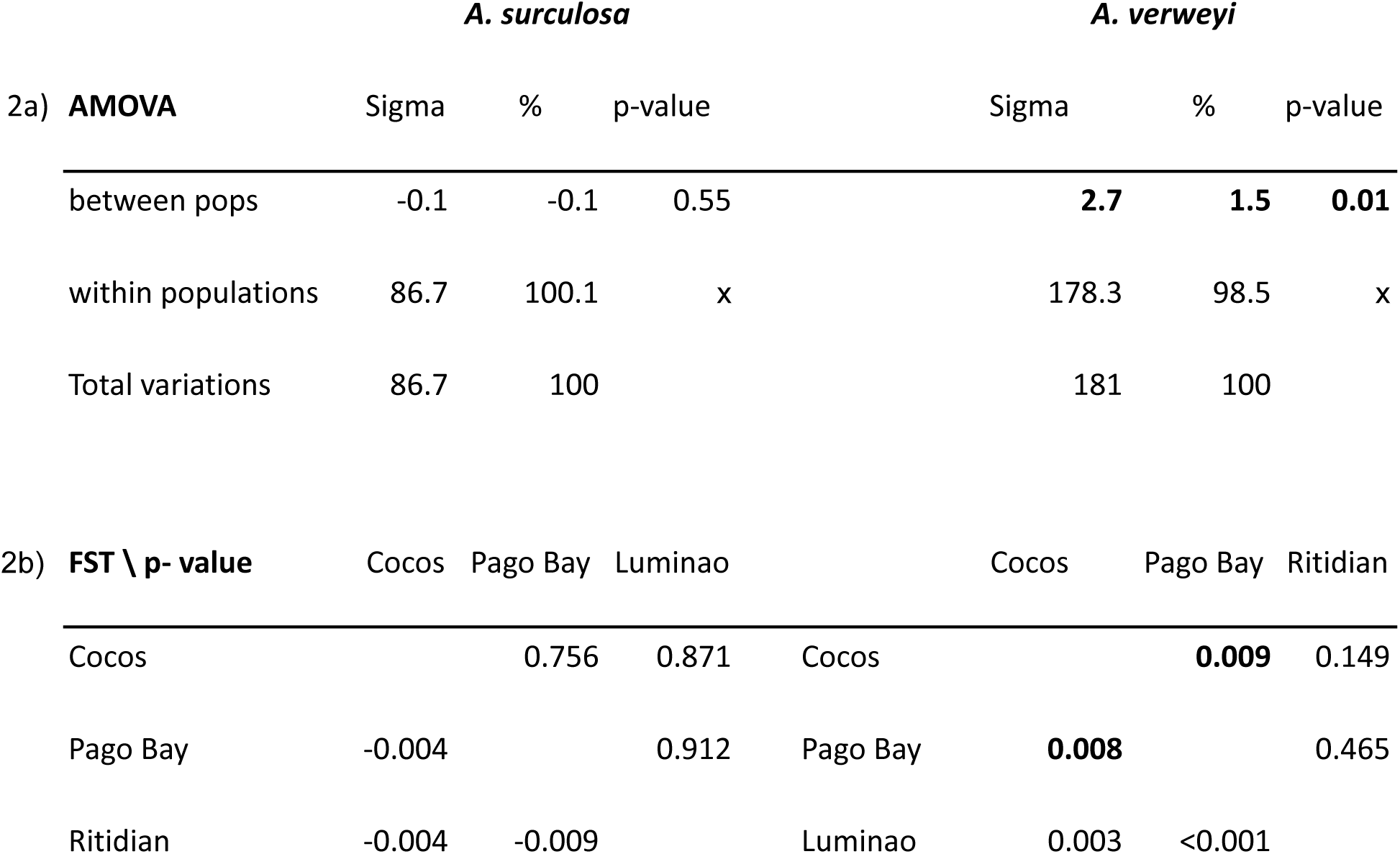
AMOVA and FST. a) Analysis of molecular variance (AMOVA) for *A. surculosa sensu* Randall & Myers (1983) (left) and *A.* cf. *verweyi* (right). b) Pairwise F_ST_ (lower diagonal) and associated p-values (upper diagonal) for *A.* cf. *verweyi* and *A*. *surculosa sensu* Randall & Myers (1983). Significance level after FDR correction = 0.027.

**Table 3.** Number of outlier SNP (upper diagonal) and their correspondent loci (putative loci under selection, lower diagonal) for *A.* cf. *verweyi* and *A. surculosa* found between populations, as detected by BayPass.

| *A. verweyi* |  |  | *A. surculosa* |  |  |  |  |
| --- | --- | --- | --- | --- | --- | --- | --- |
|  | Cocos | Pago Bay | Ritidian |  | Cocos | Pago Bay | Luminao |
| Cocos |  | 6 | 1 | Cocos |  | 1 | 2 |
| Pago Bay | 6 |  | 4 | Pago Bay | 1 |  | 1 |
| Ritidian | 1 | 5 |  | Luminao | 2 | 1 |  |

Pairwise F_ST_ values between populations are small in both species. For *A. surculosa,* all pairwise comparisons were slightly negative, i.e. zero, and insignificant (Table 2a), in line with the AMOVA results (Table 2a). In contrast, for *A.* cf. *verweyi* all three pairwise comparisons were slightly positive and one F_ST_ value was slightly elevated and significant - between Cocos Island and Pago Bay along the east side of Guam (Table 2a). This matches well with the AMOVA result of minor but significant partitioning of genetic variation between populations in *A.* cf. *verweyi* (Table 2a).

Interestingly, relatedness analyses also showed significantly different results between the two species (Figure 4): relatedness among *A. surculosa sensu* Randall & Myers (1983) samples was low overall, as expected for unrelated individuals in a sexually outcrossing population. Only 1 sample pair (out of 584 pairwise comparisons, 0.2%) had a 3rd degree relatedness.

**Fig. 4.**
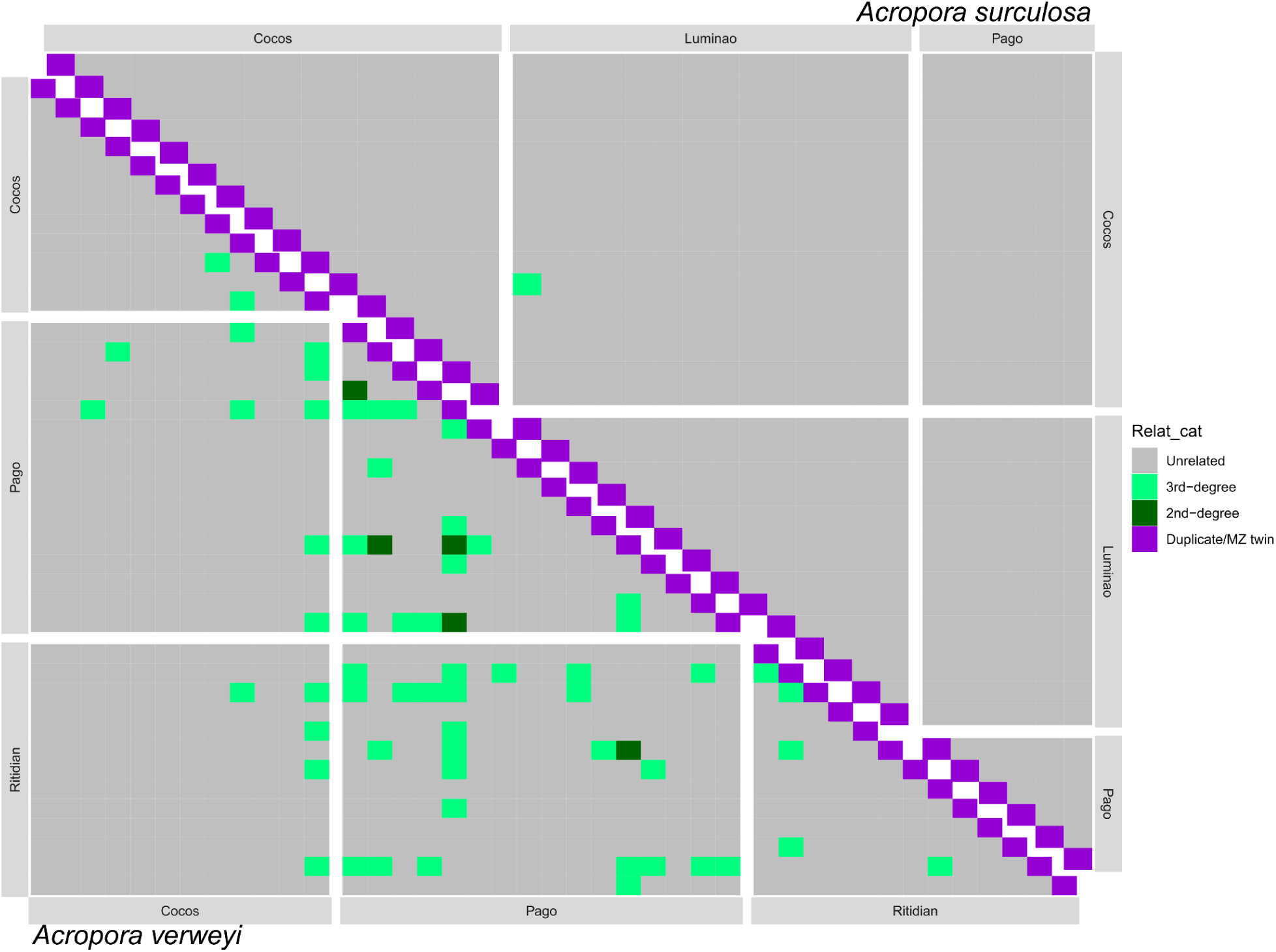
Pairwise relatedness among individuals for *A. surculosa sensu* Randall & Myers (1983) (upper diagonal) and *A.* cf. *verweyi* (lower diagonal).

In contrast, for *A.* cf. *verweyi*, 60 sample pairs (7.5% or out of 799 comparisons) had a 3rd degree relatedness and an additional 5 pairs (0.6%) had a 2nd degree relatedness. As expected, most of the 2nd degree related individuals were found together in the same population (4/5 pairs) but surprisingly, all of them were found in just one single population, Pago Bay. In addition, the only other pair of 2nd degree relatives was found between Pago Bay and Ritidian, and most of the 3rd degree relatives pairs were also found either within Pago Bay (14/60) or involving at least one colony from Pago Bay (n = 34/60; 25/60 of those between Pago and Ritidian). The number of 3rd degree relatives within Ritidian (5/60) and Cocos (2/60) as well as between them (5/60) was much lower and no second degree relatives were found within or between these populations. All together, this data indicates a high retention of larvae in Pago Bay compared to other populations and good connectivity between Ritidian and Pago bay.

In line with these results, migration analyses indicate high levels of self-seeding and local recruitment in both species (Figure 5, Table S3). For both species, one population was identified as a major source of larvae for other locations but interestingly the main source population differs between the two species: In the case of *A.* cf. *verweyi*, the main source population is Pago Bay, on the east coast of Guam, which contributes ∼25% of larval recruits to both Cocos and Ritidian. This result is in line with the observations in the relatedness analyses that found numerous relatives of Pago Bay samples in other populations (Figure 4). For *A. surculosa sensu,* Cocos, off the southern tip of Guam, was the most important source population for both Pago Bay and Luminao. Interestingly, both species-specific source populations were no significant source of larvae in the other species.

**Fig. 5.**
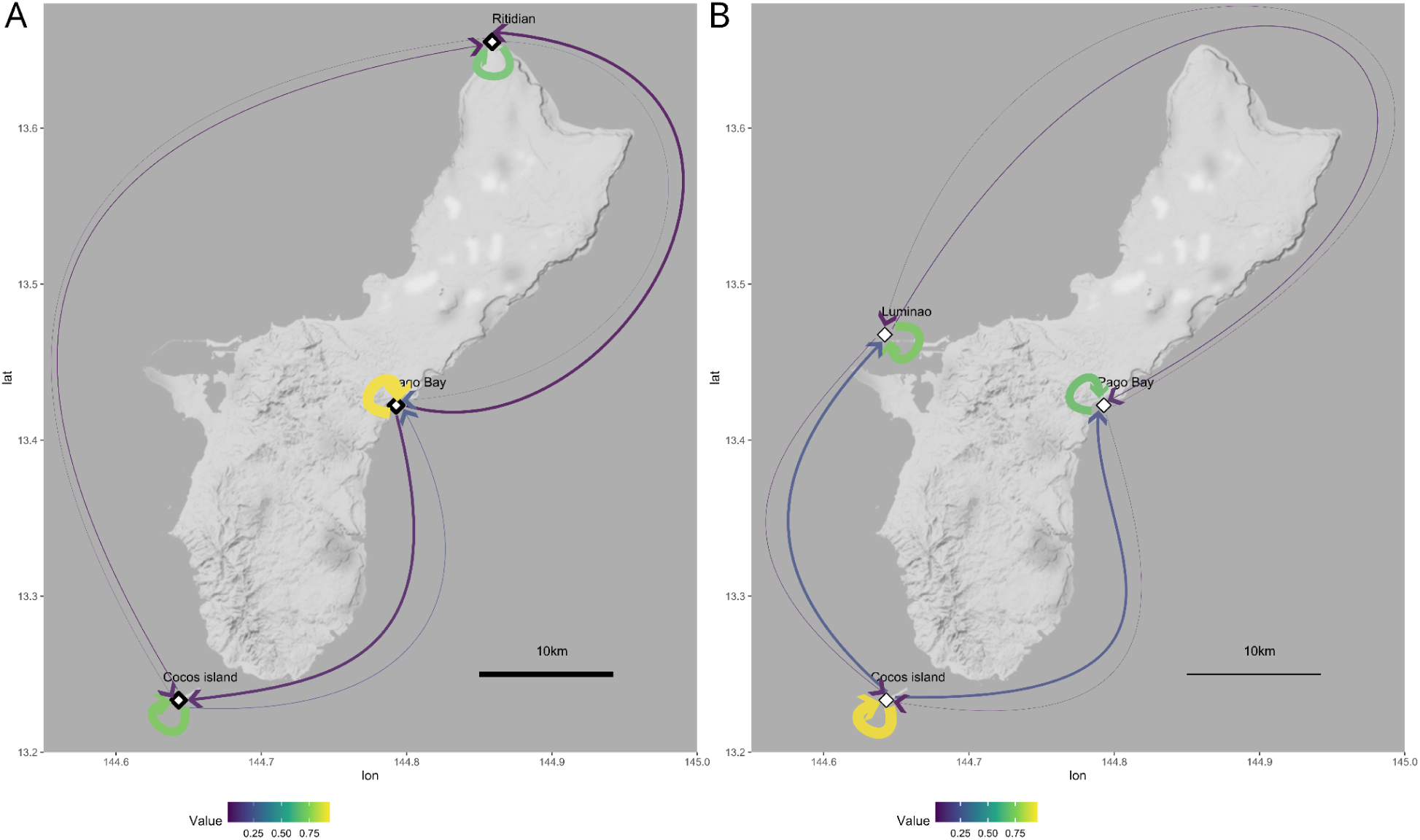
Migration estimates. Arrow color and width indicate the proportion of individuals in each population that originated in the population itself or in other populations, as calculated with BA3-SNPs for A) *A.* cf. *verweyi* and B) *A. surculosa sensu* Randall & Myers (1983). For detailed values and confidence intervals see Table S3.

Selection analysis with BayPass identified 11 putative loci under selection in *A.* cf. *verweyi* and 4 in *A. surculosa* in the three pairwise comparisons among populations - no locus was identified as an outlier in more than one pairwise comparison. Similarly, the genome-wide Tajima D analyses identified 9 loci under selection (< −2) for *A. surculosa* and 11 for *A.* cf. *verweyi*. In contrast, all SNP-based Tajima D values were higher than −2, indicating no significant signature of selection (Figure S5). Similarly, Bayescan analyses did not find any putative loci under selection for either *A.* cf. *verweyi* or *A. surculosa* (Table S3).

Finally, the most abundant Symbiodinacea genus in both species was *Cladocopium* (Figure S6). *Durusdinium* was present in all four *A. surculosa* populations and in two *A.* cf. *verweyi* samples from Cocos. In addition, there was some background presence of *Breviolum* and *Symbiodinium* in a handful of samples across populations.

## Discussion

In this study, we report comparative genomic analysis of two congeneric coral species, *Acropora surculosa sensu* Randall & Myers (1983) and *A.* cf. *verweyi* Veron & Wallace 1984, from fore reefs slopes around a small oceanic island. Phylogenomic analyses revealed that the two target *Acropora* species are only distantly related, and evolved separately since ∼50 million years (Shinzato et al. 2021) (Figure 2). Population genomic analyses indicate considerable differences in genetic and genotypic diversity, spatial genetic structure, genetic diversity, LD-based effective population sizes and the proportion of close-related individuals. These results suggest different evolutionary patterns and demographic histories between the two species, despite their apparent ecologic similarities. On the other hand, both species showed virtually no population structure around Guam and host similar photo-symbiont communities.

No major differences in the categories most expected to show indications of bleaching resilience were detected between the two species: Only a handful of putative loci under selection were identified in both species and slightly more in the more susceptible *A. verweji*. Clonality was higher in *A. verweyi*, which may have contributed to higher mortality in the more susceptible species. And the effective population size was higher in the more susceptible *A. verweyi*, which was unexpected and indicates no major impact of recent mortalities on the standing genetic diversity in *A. verweyi*.

For a more comprehensive picture, we compare and contrast our results with recent population genomic results for *A.* cf. *pulchra* (Rios et al. 2024), which is more closely related to *A. surculosa* (as discussed below) but has a branching, fragile morphology more comparable to *A.* cf. *verweyi*. Interestingly, *A.* cf. *pulchra* inhabits lagoons and back reefs, i.e. is ecologically distinct, compared to the other two fore reef target species, which creates mutually-exclusive patterns of relative similarities among two out of the three species for habitat/ecology, colony morphology and relative relatedness/phylogeny.

### Phylogenetic placement

The overall topology of our phylogenomic results is virtually identical to the most recent, comprehensive genome-based phylogeny of *Acropora* species (Shinzato et al. 2021) and other recent phylogenomic analyses of *Acropora* corals (Cowman et al. 2020; Quek et al. 2023). There are two minor differences with limited node support: First, *Acropora selago*, which formed a clade with newly added *A.* cf. *pulchra* and *A. millepora* here (node support: 100/99, Figure 2), was closely related to a clade composed of *A. cythera* and *A. hyacinthus* (node support: 89) in Shinzato et al (2021). Secondly and of no relevance for the present study, the relative position of *A. intermedia* and *A. gemmifera* is different and not well supported here (bootstrap support 56).

The two main target species of this study, *Acropora* cf. *verweyi* and *A. surculosa sensu* Randall & Myers (1983) belong to two different clades that represent the earliest divergence within the genus (Figure 2); these clades separated at least 50 million years ago, during the early Eocene (Shinzato et al. 2021). The presence of this early split in *Acropora* history is also present in many other studies, if at least one of these species (*A. verweyi*, *A. yongei* and/or *A. tenuis)* is included (Cowman et al. 2020; Quek et al. 2023).

As detailed in the introduction, *A. surculosa* (Dana, 1846) was previously synonymized with *A. hyacinthus* (Veron & Wallace 1984) but *A. surculosa/hyacinthus* from the Mariana Islands is currently being described as a separate species by Fitzgerald et al. (pers. comm.). Therefore, we have used *A. surculosa sensu* Randal 1983 to avoid confusion with other lineages of the *A. hyacinthus* species complex (Ladner & Palumbi 2012). The results presented here do not provide conclusive evidence that *A. surculosa sensu* Randal & Myers (1983) is a clearly distinct, separate species from *A. hyacinthus* (Figure 2). It is notable, though, that the branch length between *A. hyacinthus* from Okinawa and *A. surculosa* from Guam (0.0054 + 0.0035 nucleotide substitutions per site) is longer than the branch length between the clearly distinct species *A.* cf. *pulchra* and *A. millepora* (0.0020 + 0.0054) and roughly comparable to the distance between *A. tenuis* and *A. yongei* (0.0076 + 0.0065). Resolving this taxonomic issue will require further research within a broader phylogenomic context, including type specimen and/or locations and other aspects of species identity, like morphometrics, reproduction, ecology etc.

The third *Acropora* species from Guam, *A.* cf. *pulchra* (Rios et al. 2024) formed a clade with *A. millepora* from the GBR (Fuller et al. 2020) (Figure 2). The position of *A. millepora* is consistent in several recent studies (Gault et al. 2021; Huang & Roy 2015; Quek et al. 2023) but the position of *A. pulchra* has been much less consistent. For example, *A. pulchra* from Australia is placed in a different clade with *A. aspera* in a study, based on seven mitochondrial markers and morphometrics (e.g. in Gault et al. 2021; Huang & Roy 2015) Another study, based on multiple genomic datasets, placed *A. pulchra* from Australia next to a clade composed of *A. intermedia* and *A. gemmifera* - but with low node support (bootstrap = 64; (Quek et al. 2023)). In addition to methodological differences, some of these discrepancies might be due to different sampling locations and may indicate potentially cryptic *A. pulchra* species. This is in line with population genetic results that indicate extremely low genetic diversity for Guam *A.* cf. *pulchra*, which indicates a relatively small effective population size, more indicative of an island endemic than a broadly distributed species like *A. pulchra* (Rios et al. 2024).

For the three species from Guam, *A. surculosa, A.* cf. *verweyi* and *A.* cf. *pulchra,* our phylogenetic results thus indicate two rather closely related species (*A. surculosa* & *A.* cf. *pulchra*), whose lineages likely separated around 30 mya (Figure 2, Shinzato et al, 2021). In contrast, *A.* cf. *verweyi* diverged from these two roughly 50 mya (Figure 2, Shinzato et al, 2021), i.e. almost twice as long ago. This is in contrast to the habitat preferences of these three species, which indicate more ecologic similarities among the two fore reef species, *A. surculosa* and *A.* cf. *verweyi*, compared to the back reef *A.* cf. *pulchra*.

### Population Genetics

#### Population composition

*Acropora surculosa sensu* Randall & Myers (1983) and *A.* cf. *verweyi* exhibited remarkable differences in their clonality levels (Table 1/ Figure S1): while ∼20% of the 51 *A.* cf. *verweyi* colonies were clonal, i.e. non-unique genotypes, no clones were observed among the 36 *A. surculosa* samples. Importantly, samples for both species were collected only every 10 m and since *A.* cf. *verweyi* clones are spatially clustered (Figure S2), the proportion of clones is likely much higher if neighboring colonies are included. Likewise, the existence of *A. surculosa* clones cannot be excluded since clones may have been found if nearbier colonies were examined (e.g. Hämmerli & Reusch 2003). Nonetheless, there is a clear and significant (p-value = 0.0045) difference in the abundance and proportion of clones in these two species, indicating significant differences in their basic population composition and the importance of sexual vs. asexual reproduction.

The observed differences between *A. surculosa* and *A.* cf. *verweyi* can likely be attributed, at least partially, to their distinct morphologies. *Acropora hyacinthus’* growth form is frequently described as tabular (Veron 2000) but the Marianas morphospecies (*A. surculosa sensu* Randal 1983, see discussion above) is corymbose, with relatively thick and robust branches (Figure 1). Although *A.* cf. *verweyi* is frequently described as corymbose as well (Veron 2000; Wallace et al. 2012), the Marianas *A*. cf. *verweyi* forms clumps with rather thin, terete, sometimes elongate branches that break easily (Figure 1). Broken branches of *A.* cf. *verweyi* are capable of reattaching and regenerating into a new, clonal, coral colony (Knoester et al. 2019, pers. observ.), i.e. vegetative fragmentation. The fact that all clonemates were found in the same population, often in close proximity (significant ramet SGS over 10-20m; Figure S1), supports the conclusion that vegetative fragmentation is the most parsimonious explanation for the origin of these genetically identical, clonal colonies, which depends heavily on a suitable, i.e. fragile, colony morphology.

A recent study of *Acropora* cf. *pulchra* populations in Guam showed much higher levels of clonality (∼50%), using comparable sampling and analyses (Rios et al. 2024). In Guam’s waters, *Acropora* cf*. pulchra* occurs in shallow lagoons and reef flat platform environments but has a branching morphology as well, which may contribute to the elevated levels of clonality in this species. Branching morphologies, like that of *A.* cf. *pulchra* and *A.* cf. *verweyi*, likely lead to higher rates of branch breaking and more frequent vegetative fragmentation (Pipithkul et al. 2021; Wallace 1999). In addition, the generally calmer environment likely facilitates the stabilization of *A. pulchra* fragments broken during relatively infrequent high wave energy events so that they can resettle and grow into stable colonies without much wave disruption. Similar differences in clonality and genetic diversity levels have been observed in other *Acropora* species with distinct colony morphologies and habitat preference. For example, in a small-scale study in Guadeloupe, fragile back reef *A. cervicornis* were 100% clonal (i.e. one and the same genotype in all 80 samples), while robust fore reef *A. palmata* had much more genotypic diversity (10 different genotypes in 80 samples) (Japaud et al. 2015). Among robust *Acropora* species in the Indo-Pacific, clonality levels tends to be similarly low or absent, e.g. in *A. tenuis* in the Ryukyu islands (Zayasu et al. 2018) and in East Africa (Van Der Ven et al. 2016), in *A. hyacinthus* in Japan (Nakabayashi et al. 2019) or in *A. solitaryensis* on Australia (Noreen et al. 2013). In contrast, more fragile *Acropora* species like *A. pruinosa*, with a similar colony morphology as *A. verweyi*, tend to have much higher levels of clonality (e.g. Pipithkul et al. 2021). Conservation plans for species with high levels of clonality like *A. pulchra*, *A. cervicornis* or *A. verweyi* demand careful consideration when selecting individuals for restoration or sexual reproduction to ensure sufficient genotypic diversity (e.g. Koch 2021) - while such concerns are less pressing for clonal species, like *A. surculosa* or *A. palmata*. Consequently, habitat and morphology may be used as preliminary indicators for genotypic diversity until genetic results are made available.

#### Genetic Diversity and Isolation

One of the most striking aspects of our genetic diversity estimates are the significantly higher levels of SNP-based genetic diversity estimates (H_E_ & H_O_) in *A.* cf. *verweyi* but significantly higher whole genome-based parameters in *A. surculosa* (Table 1 & S2). We propose that this difference is due to a significantly higher number of rare alleles in *A. surculosa*, i.e. many heterozygous sites (and thus a high genome-based heterozygosity), but low proportions of heterozygous individuals per SNP (low SNP-based heterozygosity). This interpretation is supported by the SFS (Figure S4), which shows a much higher proportion of singletons and low frequency alleles in *A. surculosa*, while *A.* cf. *verweyi* had a much higher proportion of common SNPs (Figure S4). Together with the negative Tajima’s D values (Figure S5), this indicates a recent population expansion (Biswas & Akey 2006) of *A. surculosa* on Guam. Tajima D results are also in line with our original hypothesis of higher values in *A.* cf. *verweyi*, which we expected due to recent bleaching events that caused more mortality and thus more severe population bottlenecks in *A.* cf. *verweyi* (Raymundo et al. 2017, 2022; Reynolds et al. 2014, pers. observ.). For example, in October 2018, *A.* cf. *verweyi* had completely disappeared from one of our sampling sites (Luminao), while *A. surculosa* was still abundantly present (pers. observ.). These recent mortality events likely removed rare alleles and thus any signatures of previous population expansions in *A.* cf. *verweyi* and most likely offset any prior/actual population expansion as well. Both datasets further indicated significantly higher heterozygote deficits in *A. surculosa* (Tables 1, S1), which further supports the argument of a population expansion in this species, as expanding populations usually present excesses of rare alleles and heterozygosity relative to the number of observed alleles (Excoffier et al. 2009; Maruyama & Fuerst 1984, 1985).

Both observed and expected heterozygosity estimates for both species were slightly higher than the extremely low estimates for *A.* cf. *pulchra* on Guam (Rios et al. 2024) but lower than for most other *Acropora* populations. For example, genome-wide heterozygosity estimates for *A. hyacinthus* in Japan (∼0.0024-0.0030) (Fifer et al. 2022) are comparable to our results for *A. surculosa* (0.0039) and thus about twice as high as our estimates for *A.* cf. *verweyi* (0.0019)(Table S2). Our SNP-based heterozygosities for *A. surculosa* (0.154) estimates are most comparable to the lowest estimates for Caribbean *A. cervicornis* and *A. palmata* (0.109-0.156) (Kitchen et al. 2020) although other studies have suggested much higher heterozygosities for those two species (0.20-0.39), e.g. (Devlin-Durante & Baums 2017; García-Urueña et al. 2022), similar to some estimates for *A. tenuis* and *A. digitifera* from Okinawa (Zayasu et al. 2021), the lowest of which are more in line with our *A. verweyi* (0.250) but some much higher. This suggests that Guam’s geographical isolation may play a major role in the low heterozygosity (i.e., genetic diversity) observed in these two species, but it’s not necessarily a general pattern for all the corals in the island’s waters.

The significantly higher effective population sizes of *A.* cf. *verweyi* are curious since long-term monitoring data does not suggest a larger consensus population on Guam (Burdick et al. 2023). The most parsimonious explanation is that *A.* cf. *verweyi* is globally more abundant, which is supported, for example, by its presumably wider global distribution (although likely still smaller than indicated in (Veron et al. 2024). In contrast, recent taxonomic studies (discussed above) indicate that *A. surculosa sensu* Randal 1983 has a much narrower distribution range, potentially limited to the Mariana Islands (Fitzgerald & Bridge, pers. com.), which would align well with the effective population size estimates observed here. In line with these results, the effective population size estimate for *A. surculosa* on Guam (Ne ∼250) is notably smaller than recently reported for *A. hyacinthus* in Japan (Ne ∼10,000; Fifer et al. 2022). Although some of this discrepancy may be partially due to analytical differences (ANGSD-based multimodel inference in MOMENTS vs. STACKS-based NeEstimator), both estimates are based on RAD-Seq derived, genome-wide SNP data. The effective population size estimates for both *A. surculosa* and *A.* cf. *verweyi* on Guam are also >100-fold lower than the consensus population sizes for 15 *Acropora* species in Australia - some of them with very restricted distribution ranges (Richards et al. 2008). These estimates, however, are not based on molecular markers but based on census data only (Richards et al. 2008), which makes them difficult to compare directly. Nonetheless, both comparisons indicate highest effective population sizes elsewhere, which indicates low effective population sizes and significant isolation for two studied *Acropora* populations on Guam. For both species, Ne estimates exceed the threshold of 100 individuals required to avoid inbreeding depression but fall below the threshold of 1,000 individuals needed to maintain evolutionary potential (Frankham et al. 2014), highlighting the potential vulnerability of these populations to genetic drift and environmental changes.

#### Population structure

Both target species showed very low levels of population structure around Guam (Table 2a). This is not unexpected given their broadcast spawning reproductive strategy and the short spatial distances between sampled populations (<50km). In addition, Guam is surrounded by coral reefs, with few habitat gaps (Burdick et al., 2008), so our samples were collected from an almost continuous circular population around the island versus discrete populations with distinct spatial separations. While the migration analyses revealed that self-seeding is the predominant mechanism of recruitment for both species (Figure 5, Table S3), the main source populations we identified supply ∼20-25% of recruits to all other populations. Populations of both species are thus well connected around Guam, preventing the establishment of significant population structures (Table 2, Figure 3).

Despite the overall weak population structure, a small but significant proportion of genetic variation was partitioned between populations in *A.* cf. *verweyi* and a small but significant genetic differentiation was observed between Pago Bay and Cocos Island (Table 2). The significant differentiation in *A.* cf. *verweyi* along the east side of Guam (Figure 1) may be due to the temporally-variable current regime on the east coast of the island, where the North-Equatorial Current (NEC) arrives and splits into a northbound and a southbound coastal current (Wolanski et al. 2003). This split likely creates a barrier to gene flow, leading to the observed differentiation. Interestingly, *A. surculosa* does not appear to be affected, which could be due to a slight offset in reproduction times between these two species: in Guam *A. surculosa* has been consistently reported to spawn between the last quarter and waning crescent moon in July while *A.* cf. *verweyi* (as *A. squarrosa*) has been reported to spawn between the waxing crescent and first quarter moon in July) (Richmond & Hunter 1990; but note that Kenyon 1994 reported a single *A*. cf. *verweyi* colony spawning during the same moon phase as *A. surculosa*). These differences in reproduction times and the corresponding difference in currents (Wolanski et al. 2003) may structure these species differently but further research is necessary to explore and explain the observed differences.

The population structure of *A. surculosa* and *A.* cf. *verweyi* on Guam contrast with findings for *A.* cf. *pulchra*, which revealed significant population structure overall and significant differentiation between most population pairs. In addition, the most significant levels of differentiation were detected between north-western and south-western *A.* cf. *pulchra* populations (Rios et al. 2024). These differences in population structure are likely influenced by their distinct habitat preferences. In Guam, fore reef habitats suitable for *A.* cf. *verweyi* and *A. surculosa* are almost continuous around the island. In contrast, shallow lagoon and reef platform habitats suitable for *A.* cf. *pulchra* are much less continuous and often separated by significant gaps (Burdick et al. 2008), which likely influences the connectivity among these distinct populations. More pronounced population structure, i.e. less connectivity, in back reef compared to fore reef populations has also been observed in other studies elsewhere, for example in back reef *A. cervicornis* compared to fore reef *A. palmata* (forereef) on the Colombian caribbean coast (García-Urueña et al. 2022). Currents and water motion are generally more reduced in lagoons compared to fore reef locations (Kennedy 2002), which may slow down larval transport and thereby reduce connectivity.

The similar results of the selection analyses with BayPass (11 putative loci in *A.* cf. *verweyi*, 4 in *A. surculosa*) and the genome-wide Tajima D analyses (11 loci in *A.* cf. *verweyi,* 9 in *A. surculosa*) do support each other and are in line with the slightly more pronounced population structure of *A.* cf. *verweyi*, facilitating potential local adaptations. The fact that SNP-based Tajima D estimates and Bayescan analyses did not find any putative loci under selection indicates that the signatures of selection are not very strong or pronounced (Table S3). Similarly, (Rios et al. 2024) identified significantly more putative loci under selection in *A.* cf. *pulchra* on Guam (52 loci across 4 populations), using the same approach (BayPass) on a very similar ddRAD dataset. These differences are likely driven by the environmentally significantly more stable fore reef habitat of *A.* cf. *verweyi* and *A. surculosa* compared to the more variable back reef habitat of *A.* cf. *pulchra* (Burdick et al. 2008; Cook et al. 1990; McCabe et al. 2010), which can weaken selective forces driving divergent adaptations over small spatial scales. Moreover, the stronger population structure in *A.* cf. *pulchra* may further facilitate divergent local adaptations, as gene flow can counteract gene frequency changes due to selection, limiting local adaptation (Lenormand 2002). This aligns well with the observed decrease in putative loci under selection from the significantly structured *A.* cf. *pulchra* populations (52 loci), to the weakly structured *A.* cf. *verweyi* populations (11 loci), to the unstructured *A. surculosa* populations (4 loci).

And finally, our analyses indicate a relatively uniform composition of the symbiotic communities across both *A.* cf. *verweyi* and *A. surculosa* colonies on Guam. Both species predominantly host *Cladocopium*, with minor proportions of *Durusdinium* observed in approximately half of the *A. surculosa* colonies and a few *A.* cf. *verweyi*. While several studies have found much more variability in other *Acropora* species around small oceanic islands (Rios et al. 2024; Rouzé et al. 2017), it is common for reef coral species to exhibit a primary association with a particular clade (e.g. Berkelmans & Van Oppen 2006; Rouzé et al. 2017), and *Cladocopium* is the most dominant genera in other *Acropora* species (e.g. Davies et al. 2020; LaJeunesse et al. 2004). While our findings align with these general trends, they also underscore a concerning lack of symbiont diversity across all colonies of both species. The background presence of *Durusdinium*, which has been reported to be associated with substantially higher thermal tolerance (Berkelmans & Van Oppen 2006; Fuller et al. 2020; Klueter et al. 2017; Morikawa & Palumbi 2019), may provide some thermal resilience. However, its limited distribution within these populations might not be sufficient to confer widespread resilience, suggesting potential vulnerabilities of these coral populations to future bleaching events.

## Conclusion

In this study, we compared the population genetics of two congeneric, sympatric, fore reef coral species, *Acropora* cf. *verweyi* and *A. surculosa sensu* Randall & Myers (1983), around a small oceanic island, Guam (Micronesia). Both species had weak population structures, few signatures of selection, low levels of heterozygosity, and were dominated by a single Symbiodinaceae genus. Collectively, these findings imply that the Guam populations of both species are fairly isolated and may be particularly susceptible to environmental changes. Comparisons of these results with recently published findings for another local congeneric, *A.* cf. *pulchra*, indicate numerous similarities as well as several notable differences. Moreover, these comparisons indicate intriguing patterns relating certain population genetic characteristics to morphology and ecology, which may offer valuable insights for management decisions in the absence of species-specific studies: a) species with fragile morphologies tend to exhibit a high incidence of clones (Table 1). This pattern was observed on Guam, where clonality was much higher in thinly branching *A.* cf. *verweyi* and *A. pulchra* compared to the much more robust *A*. *surculosa*, and was supported by observation in other *Acropora* species throughout the Caribbean and Indo-Pacific (see discussion above). b) habitat type can have a significant impact on population structure. Here, the two species from the fore reef exhibit very small or insignificant population structure over small to moderate distances (< 100km). In contrast, the lagoon and reef flat platform-inhabiting *A.* cf. *pulchra* displayed significantly more structure over even shorter distances around the same island. Overall our results underscore the importance of studying the genomic signatures of coral population connectivity and structure as a prerequisite for effective conservation and restoration planning. Our current dataset is limited but still very informative and these hypotheses can easily be tested and validated in future studies. Moreover, these generalizations and simplifications can be very useful as a general rule-of-thumb for management decisions in data-poor taxa.

## Materials and methods

### Sampling

Between March and September 2017, samples of *Acropora* cf. *verweyi* and *A. surculosa sensu* Randall & Myers (1983) were collected around the island of Guam (Figure 1, Table S1) to assess patterns of genetic diversity and connectivity. For *A. surculosa*, 135 samples were collected from 4 sites at the cardinal points of the islands, i.e. north, south, east and west. For *A.* cf. *verweyi*, 95 samples were collected from 3 sites only, due to the scarcity of *A.* cf. *verweyi* colonies as a result of a recent bleaching event on the eastern reefs (Figure 1, Table S1).

At each site, transect tapes were deployed at 5m depth, following the reef contour in as straight a line as possible. Thirty samples were collected every 10m along the transect to minimize the collection of genetic clones. Colonies were photographed prior to sampling and a random branch was sampled and stored in 50ml falcon tubes. Samples were transported alive to the University of Guam Marine Laboratory where a portion of each sample was preserved in 80% Ethanol and stored at −20°C for subsequent genetic work. The remaining coral nubbins were bleached overnight to remove the tissue and the skeletons were dried to serve as voucher specimens in the UOG Biorepository (see catalog numbers in Table S1). The collection of all coral samples met Guam’s and USA’s federal regulations.

### DNA extraction and genotyping

Total genomic DNA was extracted with the DNAeasy Kit (Qiagen, Hildesheim, Germany) and the GenCatch Genomic DNA Extraction Kit (Epoch, Sugar Land, TX) using manufacturer’s protocols. DNA concentrations were measured with a Qubit 3.0 dsDNA fluorometer (Thermo Fisher Scientific Inc., Waltham, MA). To generate double-digest restriction site-associated DNA (ddRAD) libraries, we followed a modified protocol based on (Combosch et al. 2017) with several elements of the Adapterama project for high level multiplexing and PCR-duplicate removal (e.g. Glenn et al. 2019) as in (Rios et al. 2024). Briefly, extracted DNA was digested using two high-fidelity restriction enzymes, PstI and MspI, and resulting fragments were ligated to custom P1 and P2 adaptors with sample-specific barcodes and primer annealing sites. DNA concentrations per sample were measured again and equal amounts of barcoded fragments of eight different samples were pooled and size-selected (320-420bp) with an E-Gel Size Select II Agarose Gel (Thermo Fisher Scientific Inc., Waltham, MA). Size-selected fragment pools were PCR-amplified, using primers with additional sequence indices and Q5 high-fidelity polymerase (New England Biolabs, Ipswich, MA). For PCR amplifications, between 15 and 25 PCR cycles (95°C for 30 s, 65°C for 30 s, 72°C for 60 s, with an initial denaturation step at 98°C for 30 s, and a final extension step at 72°C for 5 min) were used, depending on the concentrations of the resulting library pools. Between two and six separate PCR amplifications were set up per library pool and mixed subsequently to increase the diversity of sequencing pools. Sequencing pools were cleaned to remove leftover adapters and primers using Agilent beads (Agilent Technologies, Santa Clara, CA). DNA concentrations were measured again using the Qubit 3.0 and the quality of a subset of libraries was assessed using an Agilent Bioanalyzer 2100 (Agilent Technologies, Santa Clara, CA). Finally, sequencing pools were in-house single-end sequenced (150 bp) on an Illumina NextSeq500 (New England Biolabs, Ipswich, MA) at the University of Guam Marine Laboratory, following manufacturers protocols. For a subset of samples, duplicated ddRAD libraries were generated to serve as technical replicates for downstream analysis.

### Data curation and loci assembly

Raw Illumina sequences from all samples were quality trimmed to 100bp. Resulting reads were aligned to closely related *Acropora* genomes with Bowtie2 v2.3.4 (Langmead & Salzberg 2012), using --score-min L,16,1 --local -L 16. To identify the most suitable genome, a two step process was applied: At first, all raw reads were mapped to all 16 publicly available *Acropora* genomes (Fuller et al. 2020; Shinzato et al. 2021) to identify the genome with the highest proportion of mapped reads. For *A.* cf. *verweyi,* most reads mapped to the genome of *Acropora yongei* Veron & Wallace, 1984 and for *A. surculosa,* most reads mapped to the genome of *A. hyacinthus* (Dana, 1846)(both genomes from Shinzato et al. 2021). This first step identified the most “mappable” genomes but not necessarily the most closely related genome for loci assembly. In the next step, we therefore identified suitable loci for phylogenomic analyses to identify the most closely related genome for each species, as described in the following section.

### Phylogenetic analysis

For each species, aligned reads were converted to bam files and sorted using SAMtools (Li et al. 2009). Individuals with less than 50,000 aligned reads were removed from further analyses. Resulting files were used for genotyping with STACKS vs 2.59 software (Catchen et al. 2013). Individual genotypes were exported as VCF files, keeping only the first SNP of each locus and additional filters were then applied using VCFtools vs 1.13 (Danecek et al. 2011) as follows: 1. Individual loci with a depth below 5x were removed, 2. loci with a missingness value higher than 50% were removed and 3. only loci with a major allele frequency equal or higher than 0.95 (i.e. virtually monomorphic) were identified by the function ‘isPoly’ from the package ‘adegenet’ (Jombart 2008), R version 4.1.1 (R Core Team 2021), and used for phylogenetic analyses, to limit the inclusion of loci that are polymorphic within species.

Using Blast blastn v2.10.0+ (Camacho et al. 2009), the loci obtained from our two species and data for *A.* cf. *pulchra* from (Rios et al. 2024) generated with the same approach were then extracted from 16 published *Acropora* genomes, predominantly found in Shinzato et al (2021): *Acropora acuminata*, *A. awi*, *A. cytherea*, *A. digitifera*, *A. echinata*, *A. florida*, *A. gemmifera*, *A. hyacinthus*, *A. intermedia*, *A. microphthalma*, *A. muricata*, *A. nasta*, *A. selago*, *A. tenuis*, *A. yongei*, but also the *A. millepora* genome (Fuller et al. 2020). In addition, we extracted the same loci from three genomes of closely-related species in other genera as outgroups: *Astreopora myriophthalma*, *Montipora efflorescens* and *M. cactus* (Shinzato et al. 2021). Loci-specific sequences were then aligned using MEGA11 (Tamura et al. 2021) and concatenated using Phyutility v2.7.1 (Smith & Dunn 2008). Phylogenetic trees were constructed using IQ-TREE (Nguyen et al. 2015) with 10,000 bootstrap repetitions and default parameter settings, including the implemented model selection using ModelFinder. For tree plotting and annotation we used FigTree v1.4.4 (*FigTree*, 2018) and Inkscape (inkscape.org).

### Population genomic analyses

For population genomic analyses, raw reads were aligned to the most closely related *Acropora* genome, as identified before, and aligned reads were converted to bam files and sorted using SAMtools (Li et al. 2009). Samples with less than 50,000 aligned reads were removed and remaining samples were genotyped with STACKS vs 2.59 (Catchen et al. 2013). Individual genotypes were exported as VCF files but this time, only the first SNP of each locus was used to avoid SNPs with high linkage disequilibrium. VCFtools vs 1.13 was used again to filter out individual loci with a depth below 5x or extraordinarily high mean depth across all individuals (above 1.5 times the interquartile range from the dataset). Loci with a missingness value higher than 30% were also removed from the dataset. Finally, loci with a major allele frequency equal or higher than 0.95 were identified as described above and removed from the vcf file using VCFtools.

The presence of genetic clones caused by asexual reproduction was assessed by creating an Identity By State (IBS) matrix with the R SNPrelate package (Zheng et al. 2012). This similarity matrix was converted to a dissimilarity distance matrix and used to construct a hierarchical clustering tree using the R function hclust in SNPrelate. Clonality thresholds were determined by comparing technical replicates and by binned gap analysis (Figure S1). From each group of putative clones, only the best sequenced individual was retained in subsequent analyses. A Fisher’s exact test (Fisher 1935), with fisher.test function in R, was used to assess if the proportion of clones was significantly different between species.

For each species, overall observed (H_o_) and expected (H_e_) heterozygosities, as well as inbreeding coefficients (Gis) were obtained using Genodive (Meirmans & Van Tienderen 2004) based on variable SNP-positions only (“SNP-based”) and based on all positions (“genome-wide”). These indices were calculated for two different datasets, since all-position estimates are preferable because they are sample-size-independent (Schmidt et al. 2021) but SNP-based estimates are more widely available (pers. observ.). Since filtering monomorphic positions by read coverage in vcf files was not possible, none of the previous filters was applied. Instead, the alpha threshold for discovering SNPs in STACKS was lowered from 0.05 to 0.01 and the STACKS populations program was used to retain only loci that were present in 50% of all samples. To detect signatures of past demographic changes caused by differences in bleaching susceptibility, site frequency spectrums (SFS) were generated using vcf2sfs in R (Liu et al. 2018) and Tajima’s D values (Tajima 1989) were calculated in windows of 1000bp using vcftools for both datasets (i.e. genome-wide and SNP-based). Tajima’s D value over +2 and under −2 were considered significant. In addition to indicating signatures of selection, Tajima’s provides broader demographic and evolutionary context. A positive Tajima’s D indicates an absence of rare, low-frequency alleles, suggestive of a recent population expansion.

Spatial Genetic Structure (SGS) was estimated using Loiselle’s kinship coefficient (Loiselle et al. 1995) in the program SPAGeDi 1.5 (Hardy & Vekemans 2002). SGS was estimated for each of the three regions separately and for all populations combined in 10m intervals up to 200m. For *A. verweyi,* a dataset with and without clones was used to assess the spatial structure of clones vs. sexual recruits. No clones were detected in *A. surculosa*, thus only one dataset was used for SGS. 95% confidence intervals and standard errors were estimated by 10,000 permutations of the genetic and the spatial datasets. The Sp statistic (Vekemans & Hardy 2004) was calculated using the function SpSummary in the R package rSpagedi (Browne 2019). The genetic patch size was obtained as the distance corresponding to the first x-intercept of the kinship correlogram (Verity & Nichols 2014).

Effective population sizes (Ne) were calculated using the linkage disequilibrium method included in NeEstimator 2.1 (Do et al. 2014). Additionally, pairwise relatedness among individuals was calculated for both species using the function relatedness2 in VCFtools vs 1.13 (Danecek et al. 2011), which applies the method from (Manichaikul et al. 2010). We defined the degree of relationship according to the following ranges of estimated kinship coefficients: >0.354 corresponds to a mono-zygotic twin, 0.177 - 0.354 to 1st-degree relationships, 0.0884 - 0.177 to 2nd-degree relationships and 0.0442 - 0.0884 to 3rd-degree relationships. These ranges were obtained by following the KING tutorial (available at https://www.kingrelatedness.com/manual.shtml), the first software where this relationship inference algorithm was implemented (Manichaikul et al. 2010)

In order to assess population structures, pairwise *F*_ST_ values and their significance (computed by 999 permutations) were calculated with the ‘hierfstat’ package. A false discovery rate (FDR) correction for multiple comparisons was applied in all required analyses to estimate the threshold of differentiation (Narum 2006). In addition, a Discriminant Analysis of Principal Components (DAPC) was performed retaining a number of PCAs equal to one third of the number of individuals with the R software package ‘adegenet’. To assess the partitioning of genetic diversity among populations, a hierarchical Molecular Analysis of Variance (AMOVA) was conducted using the package ‘poppr’ version 2.9.3 (Kamvar et al. 2014, 2015). As this analysis is sensitive to missing data, missing genotypes were imputed with the most common genotype for that allele using R. Furthermore, an admixture analysis was performed using the software ADMIXTURE 1.3.0 (Alexander et al. 2009) and represented with the R packages ‘ggplot2’ and ‘admixture’ (Garnier 2018).

Migration among localities was estimated with BA3-SNPs (Mussmann et al. 2019; Wilson & Rannala 2003). Different numbers of iterations were tested to ensure their convergence, and finally set to 4,000,000 MCMC iterations and 1,000,000 burn-in, with a sampling interval of 100. In order to obtain an acceptance rate between 20 - 60 % as recommended by BA3-SNPs manual. Mixing parameters (migration rates dM, allele frequencies dA, and inbreeding coefficients dF) were also tested separately for each species, resulting in the following final parameters settings: dM=0.5, dA=0.85, dF=0.15 for *A. verweyi* and dM=0.5, dA=1, dF=0.2 for *A. surculosa*.

For selection analysis, we made a new dataset, following all the filters detailed above for population genetics but keeping all the SNPs per locus, as these analyses are not biased by linkage disequilibrium and having all SNPs can help to identify putative loci under selection. Finally, in order to identify outlier loci that could indicate natural selection processes between locations we used BayeScan vs 2.1 (Foll & Gaggiotti 2008). To minimize false positives, 100,000 simulations were run, specifying prior odds of 10,000 (Lotterhos & Whitlock 2014). Then, only the loci with a q-value below 0.05 were considered statistically significant outliers. In addition, we converted our vcf file to BayPass format using the reshaper_baypass script developed by Yann Dorant (gitlab.com/YDorant/Toolbox) and ran BayPass Version 2.4 (Gautier 2015). BayPass was run once to obtain the covariance matrix between populations (mat_omega). Then, this mat_omega was used as covariable to control for population structure in a set of 5 independent runs with different seeds, from which we took the median value of XtX for each SNP. Additionally, we simulated a neutral distribution of 1000 loci using the simulate.baypass function in the BayPass R script baypass_utils.R and generated 5 independent runs with it (with the same approach as above) to obtain their distribution and define the threshold to consider a locus as outlier.

#### Symbiodiniaceae genera determination

In order to detect the presence and relative abundance of different *Symbiodiniaceae* genera, we followed the method developed by Barfield et al. (2018) with small modifications. Quality filtered and trimmed ddRAD reads were mapped to a database containing the transcriptomes of *Symbiodinium, Durusdinium, Cladocopium,* and *Breviolum* with Bowtie2 v2.3.5 using the same settings described above for the coral alignment. The *Symbiodinium* and *Breviolum* transcriptomes were acquired from Bayer et al. (2012), and *Cladocopium* and *Durusdinium* transcriptomes were from Ladner and Palumbi (2012). Resulting SAM files were then used to calculate relative proportions of reads with highly unique matches (mapping quality of 40 or higher) to each *Symbiodiniaceae* transcriptome, using the perl script zooxType.pl (https://github.com/z0on/).

The use of ddRAD data for Symbiodiniaceae genera identification has been tested in previous studies (e.g. Barfield et al. 2018; Rios et al. 2024), successfully demonstrating that data obtained this way is as reliable as data obtained through ITS metabarcoding (Rios et al. 2024). However, while our approach can reliably identify the dominant symbiont genera, the presence and abundance of the minor symbionts is less accurate. We did therefore focus our discussion on those dominant genera.

## Acknowledgments

We would like to thank Hannah Weigand for providing the demultiplexing python3 script, as well as John Peralta and Jason Miller for assisting with sample collections.

All other populations were sampled under a scientific collection permit issued by the Guam Department of Agriculture to the Marine Laboratory at the University of Guam.

## Data Availability Statement

Raw read data from all individuals will be available from an NCBI SRA Bioproject upon acceptance.

## Author contributions

D.C. and S.L. conceived and designed the study. D.C., S.L., B.B., D.B., D.R. and K.P. collected samples. D.C., D.R. and K.P. generated the ddRAD data. H.T. conducted the data analysis with input from D.C.. H.T. and D.C. wrote the manuscript and all authors revised and edited the final version.

