## Supplementary material for "Evolutionary genomics of two co-occurring congeneric fore reef coral species on Guam (Mariana Islands)"

**Table S1.** Sampling information. For each species, the number of individuals sampled (Samples), the amount of samples passing the filters for coral population genetics (After filtering - host) and algal symbiont characterization (After filtering - Symbiodinacea) analysis, and the catalog number for Guam's Marine Laboratory Biorepository are provided. The last column indicates the GPS coordinates for each sampling point.

| Locality | *A. surculosa* | | | | *A. verweyi* | | | | GPS |
|---|---|---|---|---|---|---|---|---|---|
| | Samples | After filtering - host | After filtering - Symbiodinacea | Catalog numbers | Samples | After filtering - host | After filtering - Symbiodinacea | Catalog numbers | |
| Cocos Island | 42 | 16 | 25 | 288-319, 412-442 | 14 | 14 | 9 | 256-287 | 144.64; 13.23 |
| Luminao | 36 | 14 | 25 | 384-411 | 0 | 0 | 0 | | 144.64; 13.47 |
| Pago Bay | 30 | 6 | 15 | 352-383 | 22 | 22 | 19 | 320-351 | 144.79; 13.42 |
| Ritidian | 27 | **0** | 6 | 226-255 | 16 | 15 | 15 | 203-225 | 144.86; 13.66 |
| Total | 135 | 36 | 71 | | 52 | 51 | 43 | | |

**Table S2** Overall population genetic stats for both species based on all the positions sequenced.

| Species | $H_O$- Overall | $H_E$- Overall | $G_{IS}$- Overall (95% CI) |
|---|---|---|---|
| *A. verweyi* | 0.0018 (0.00175 - 0.00183) | 0.0019 (0.00190 - 0.00197) | 0.076 (0.037 - 0.085) |
| *A. surculosa* | 0.0025 (0.00249 - 0.00258) | 0.0039 (0.00384 - 0.00396) | 0.350 (0.344 - 0.356) |

**Table S3** Migration estimates. Proportion of individuals from each sampled site (rows) with origin (columns) in each of them, calculated with BA3-SNPs. Values in brackets indicate 95% confidence interval.

| ***A. verweyi*** | Origin: | | |
|---|---|---|---|
| Destination: | Cocos | **Pago Bay** | Ritidian |
| Cocos | 0.74 | **0.24** | 0.02 |
| | (0.67- 0.81) | (0.16- 0.31) | (-0.02- 0.06) |
| Pago Bay | 0.05 | **0.93** | 0.02 |
| | (0- 0.11) | (0.87- 0.99) | (-0.02- 0.05) |
| Ritidian | 0.06 | **0.25** | 0.69 |
| | (0- 0.12) | (0.18- 0.32) | (0.65- 0.73) |

| ***A. surculosa*** | Origin: | | |
|---|---|---|---|
| Destination: | **Cocos** | Pago Bay | Luminao |
| Cocos | **0.95** | 0.02 | 0.04 |
| | (0.89- 1) | (-0.02- 0.05) | (-0.01- 0.08) |
| Pago Bay | **0.22** | 0.7 | 0.07 |
| | (0.12- 0.32) | (0.64- 0.77) | (-0.01- 0.16) |
| Luminao | **0.24** | 0.02 | 0.75 |
| | (0.17- 0.31) | (-0.02- 0.06) | (0.68- 0.81) |

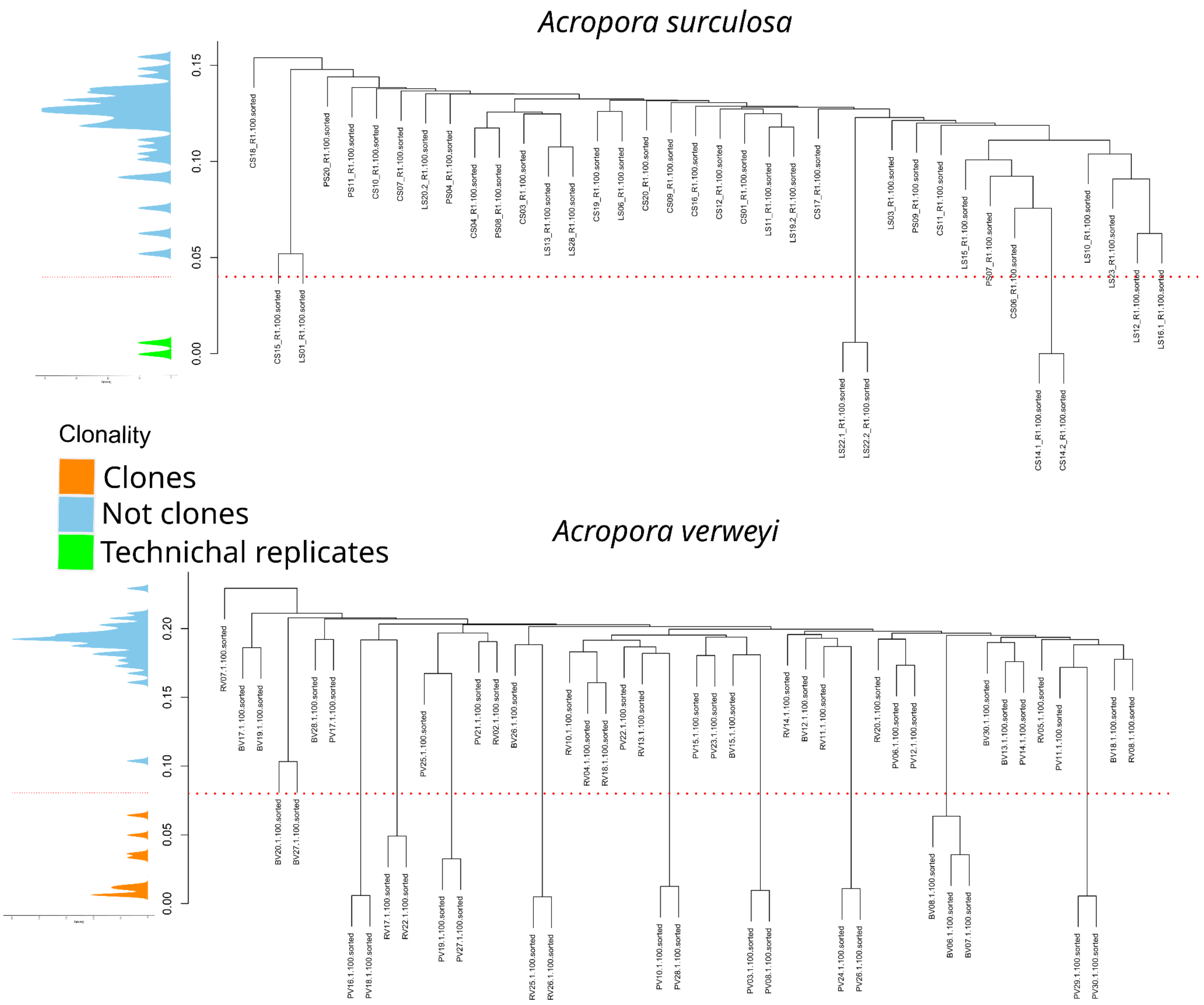


**Fig. S1** IBS-based distance trees for genetic clone filtering for *Acropora surculosa* (top) and *A. verweyi* (bottom). Red line indicates the threshold between unique genotypes and putative clones. Density plots on the left indicate density of nodes at each value.

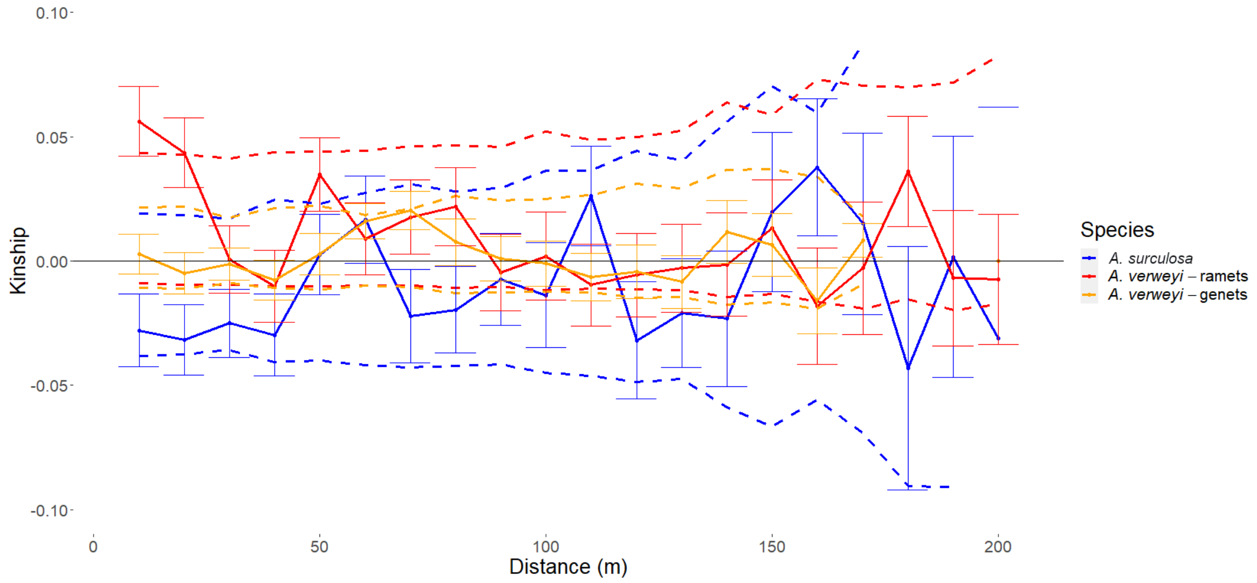


**Fig. S2** Spatial Genetic Structure (SGS). Average pairwise kinship (Loiselle et al 1995) per distance interval (10m), for both species. In the case of *A. verweyi*, analysis was performed twice, with the complete ramet dataset, including clones, and with the genet dataset used for population genetics analysis, excluding clones (see table S1). Error bars represent the SD values of each distance interval. Due to the removal of clones, the *A. verweyi* genet dataset has no values for distances greater than 170m.

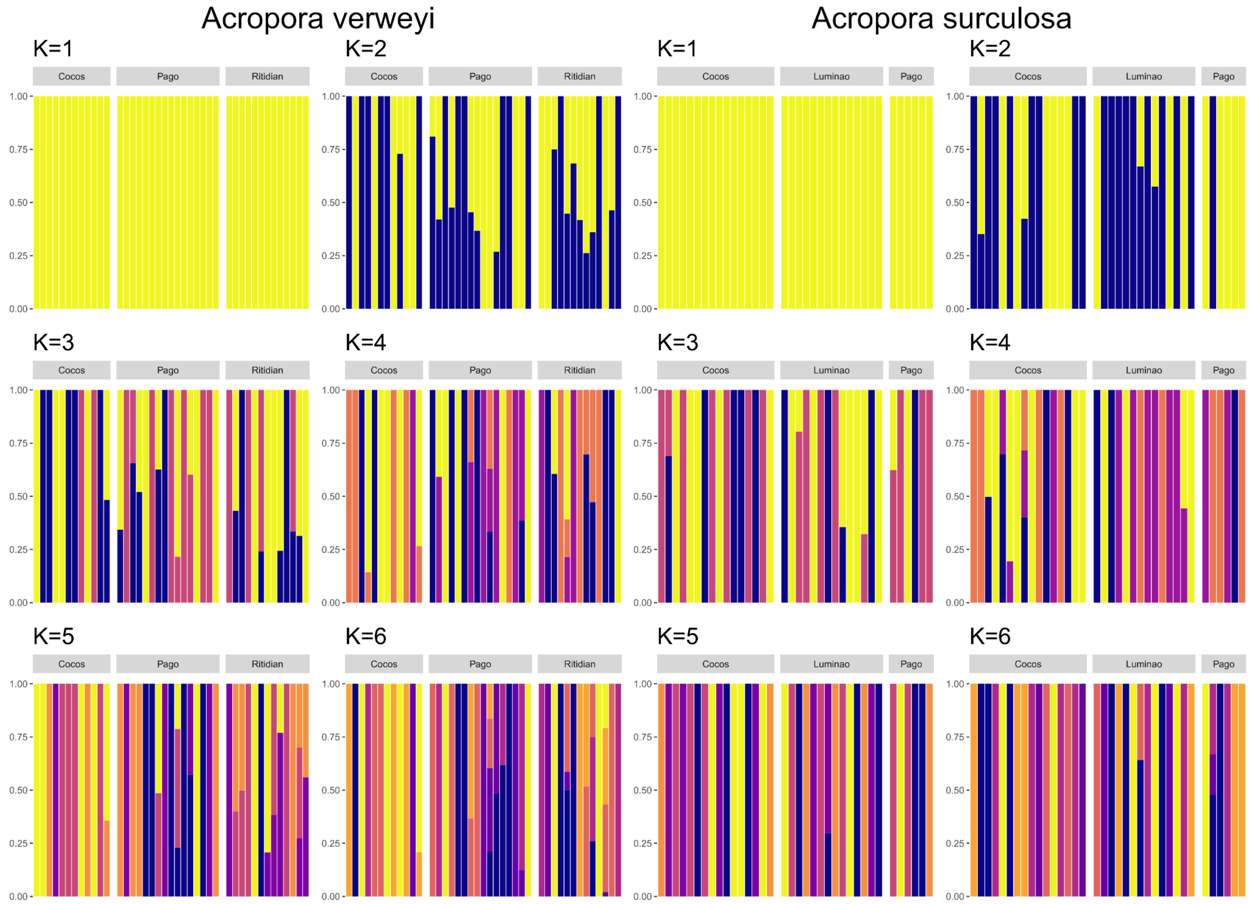


**Fig. S3** ADMIXTURE results for *Acropora verweyi* (left) and *A. surculosa* (right). Samples within localities are disposed geographically, following the sampling strategy (see Methods). The slight population structure, indicated by the AMOVA analysis, in *A. verweyi* is not readily observable. The significant pairwise differentiation between its Cocos and Ritidian populations is not very pronounced either but most notable in the K=3 and higher plots.

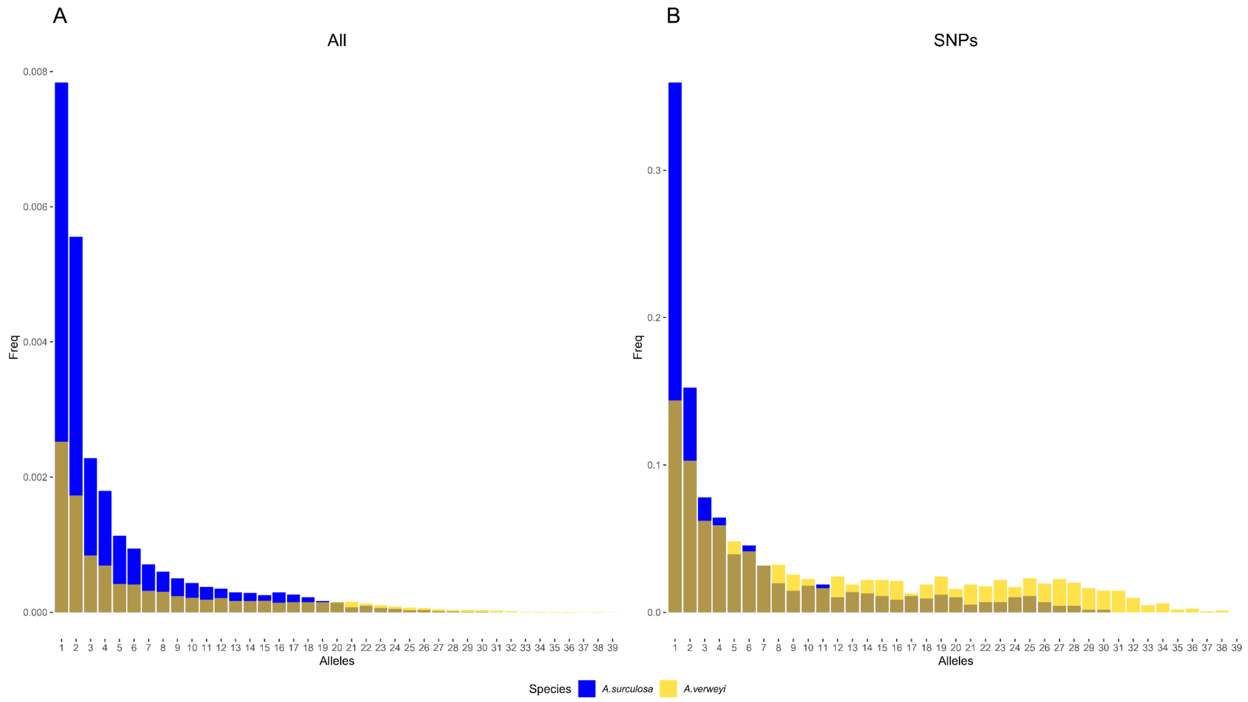


**Fig. S4** Site frequency spectrum (SFS) for *A. surculosa* (blue) and *A. verweyi* (gold), for genome-wide (A) and filtered SNPs (B) sites. Monomorphic loci were present in the genome-wide SFS (A) but have been removed here to enable a better visualization of the patterns.

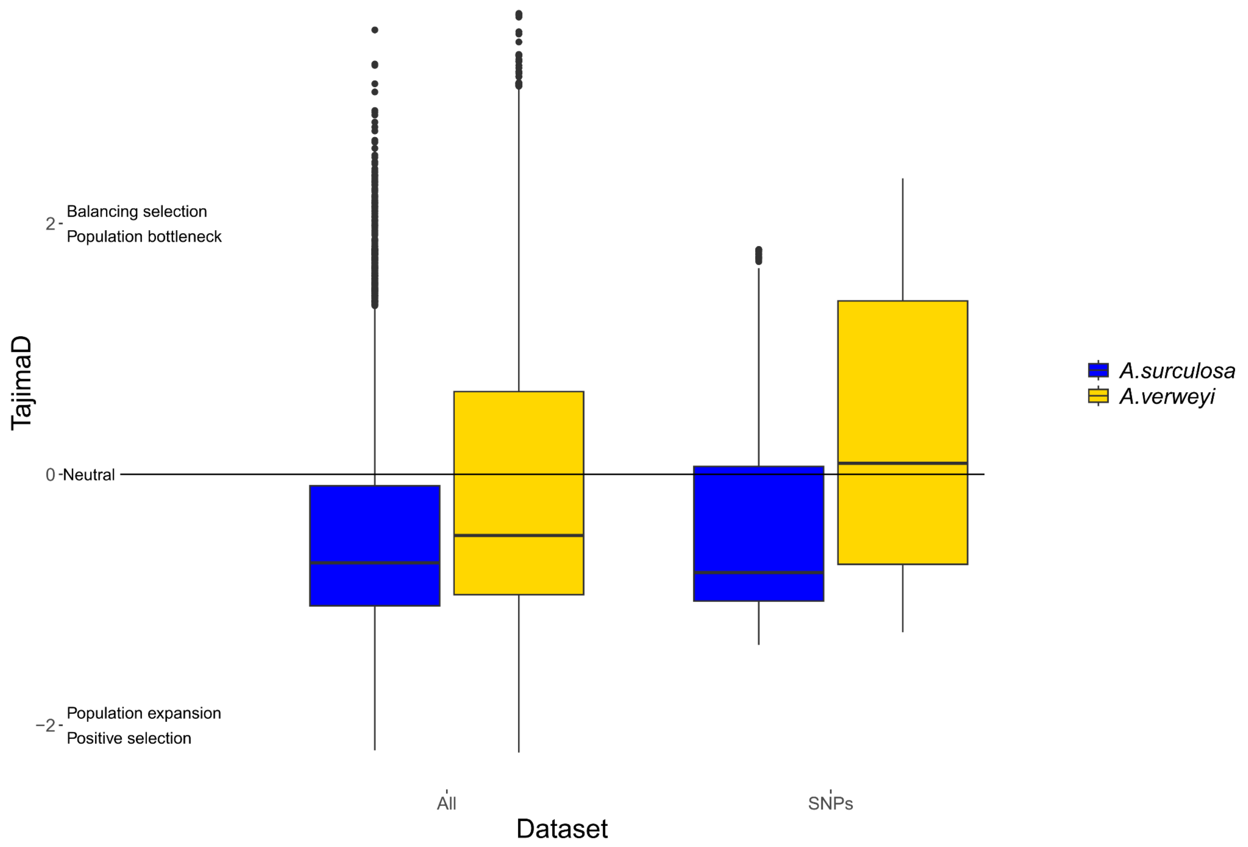


**Fig. S5** Tajima's D distribution. Tajima's D values for genome-wide (left) and filtered SNPs (right), in *A. surculosa* (blue) and *A. verweyi* (yellow). Average values±sd are -0.103±1.106 for *A. verweyi* and -0.512±0.819 for *A. surculosa* for genome-wide and 0.287±1.035 for *A. verweyi* and -0.318±0.873 for *A. surculosa* for SNP-based.

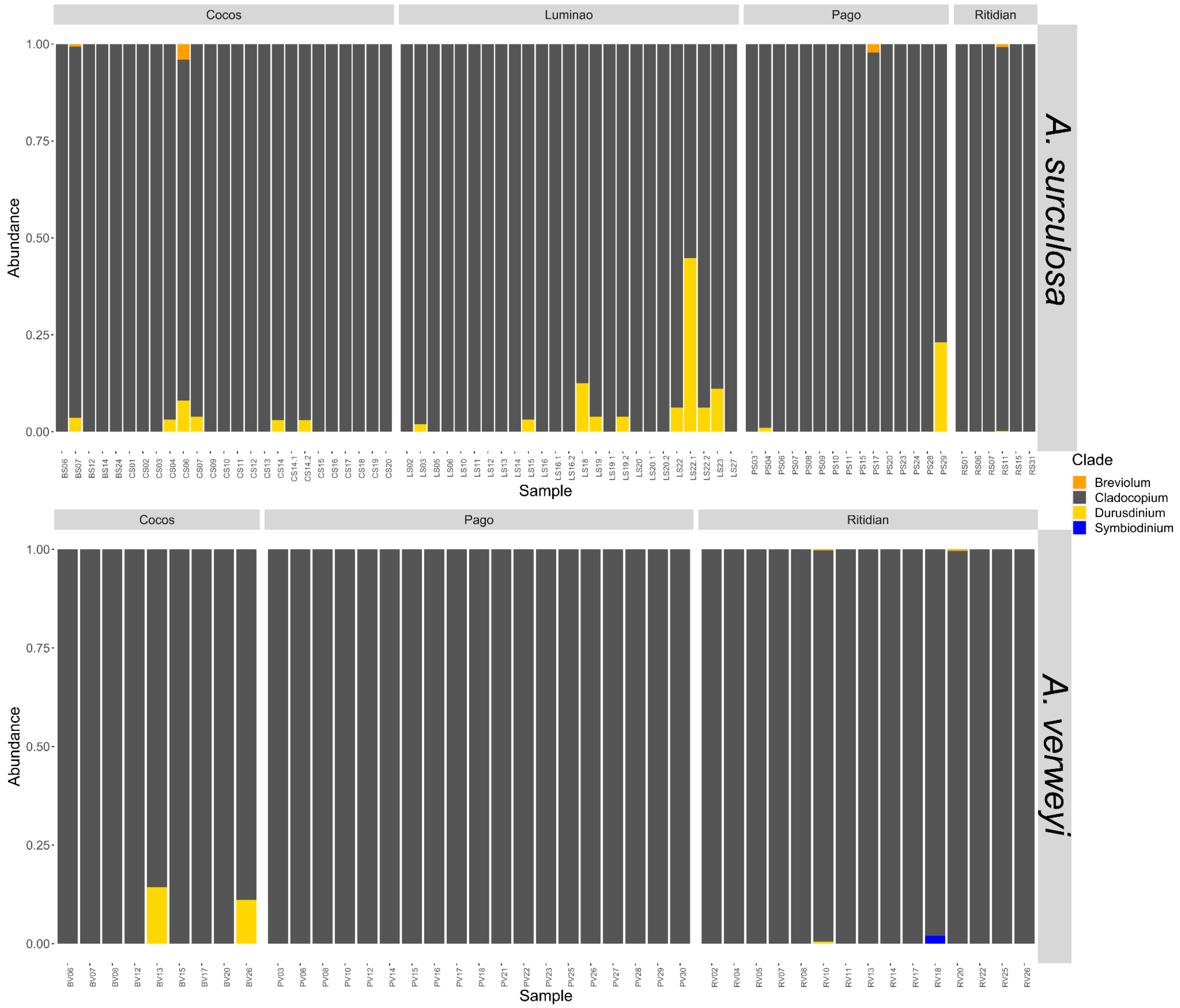


**Fig. S6** Symbiont composition. Proportion of ddRAD reads matching to transcriptomes of four different genera of algal symbionts, *Breviolum* (orange), *Cladocopium* (dark gray), *Durusdinium* (orange), and *Symbiodinium* (orange).
